# Genomic Selection Pressure Discovery using Site-Frequency Spectrum & Reduced Local-Variability Statistics in Pakistani Dera-Din-Panah Goat

**DOI:** 10.1101/2022.10.31.514560

**Authors:** Rashid Saif, Tania Mahmood, Aniqa Ejaz, Saeeda Zia

## Abstract

Population geneticists have long sought to comprehend various selection traces present in the goat genome due to natural or human-driven selection and breeding practices. As a step forward to pinpoint the selection signals in the Pakistani Dera-Din-Panah (DDP) goat breed, whole-genome pooled-sequencing (n=12) was performed and 618,236,192 clean paired-end reads were mapped against ARS1 reference goat assembly. Five different selection signal statistics were applied here using four Site-Frequency Spectrum (SFS) methods (Tajima’s D (*TD*), Fay & Wu’s H (*H*), Zeng’s E (*E*), *Pool – HMM*) and one Reduced Local-Variability approach named pooled-heterozygosity (*Hp*). The under-selection regions were annotated with significant threshold values of –*ZTD*≥4.7, –*ZH*≥6, –*ZE*≥2.5, Pool-HMM≥12, and –*ZHp*≥5, which resulted in accumulative 364 candidate gene hits, while the highest signals were observed on Chr. 4, 6, 10, 12, 15, 16, 18, 20, 27 harbor *ADAMTS6, CWC27* genes associated with body-height, *RELN, MYCBP2, FGF14, STIM1, CFAP74, GNB1, CALML6, TMEM52, FAM149A, NADK, MMP23B, OPN3* with body-weight/meat production, *FH, MFHAS1, KLKB1* with milk production, *RRM1, KMO, SPEF2, F11* with fecundity rate/reproduction, *ATP8B4* with immunity, *KIT, KMO* with pigmentation, *ERI1* with olfaction and *RHOG* with wool production traits. Furthermore, we accentuate to highlight the putative windows that were captured commonly by any of the five statistical methods applied which harbor meat production, immunity and reproduction-associated genes validating the genotype-phenotype relationship of aforementioned traits private to this goat breed. Current insight into the genomic architecture of DDP goat provides a better understanding to improve its genetic potential and other vested traits of large body size and fiber production by updating the breeding strategies to boost the livestock-based agricultural economy of the country.

## Introduction

Goats have long been domesticated since Neolithic period, when human practices shifted from hunting to farming (1). During the geographical expansion, goats have spread to diverse environments where they successfully adapted to agro-climate conditions and developed variable traits in their local surroundings (2). Indigenous breeds have characteristics which are locality-unique, with substantial regional diversity. This diversification in the phenotypic characteristics and genetic footprints in the goat genome are indications of long-term artificial selection pressures. Both natural and artificial selection lead to fixation of variants and haplotype structures that resulted in the evolution of distinct breeds within different populations (3–5).

The Dera-Din-Panah (DDP) goat breed of Pakistan is under natural selection for meat, milk and fiber production traits, however, it has also been directed towards sustainable selective breeding programme mainly to develop it as meat-cum-dairy breed (6). To identify the regions in the genome that are under selection pressure, several diversity indices including site-frequency spectrum (SFS), linkage disequilibrium, reduced local-variability, single site differentiation and haplotype-based differentiation have demonstrated to be compelling strategies (7, 8). Once selection footprints across the whole genome are identified, the causative variants contributing to subject phenotypes can be more readily recognized. In order to decipher selection signatures at a tremendously low cost, a whole-genome sequencing methodology e.g., pooled DNA sequencing (pool-seq) is now extensively being used (9). This requires maximum 20 individuals that can be pooled into one sample and a single DNA library is constructed without any sequencing biases (10). Similarly, the population genetics inferences made from pool-seq data with minimum pool size of 12 individuals is shown to be accurate, reliable and robust (9).

In the current study, finding genomic selection pressure regions in the DDP goat breed from Pakistan was aimed using four different *SFS* statistical approaches namely Tajima’s D (*TD*), Fay and Wu’s H (*H*), Zeng’s E (*E*) and Pooled Hidden Markov Model (*Pool – HMM*), while the fifth one is Pooled-Heterozygosity (*Hp*) which comes under reduced local-variability. Further annotation of the under-selection regions was also conducted to provide the list of candidate genes and associated phenotypes in this valued goat breed.

## Materials and Methods

### Animals, samples and pooled sequencing

In the current study, whole blood samples for each individual of DDP goat (Punjab/Pakistan) and Bezoar goat (Switzerland) were collected and their genomic DNA was extracted. TIANGEN biotech (Beijing) CO., Ltd. kit was utilized to extract DNA from DDP breed, while Bezoar’s DNA extraction and sequencing was carried out at Institute of Genetics, University of Bern, Switzerland. Equimolar ratios of samples were pooled and sequenced using Illumina HiSeq3000 platform that generated paired-end data of (150×2) bp of ~300mio reads per pool. The data can be accessed via ENA database under project ID: PRJEB23815. San Clemente (ARS1) reference goat genome was retrieved from NCBI. Particular features of animals used in the study are briefly described in (Table 1)

**Table 1.**
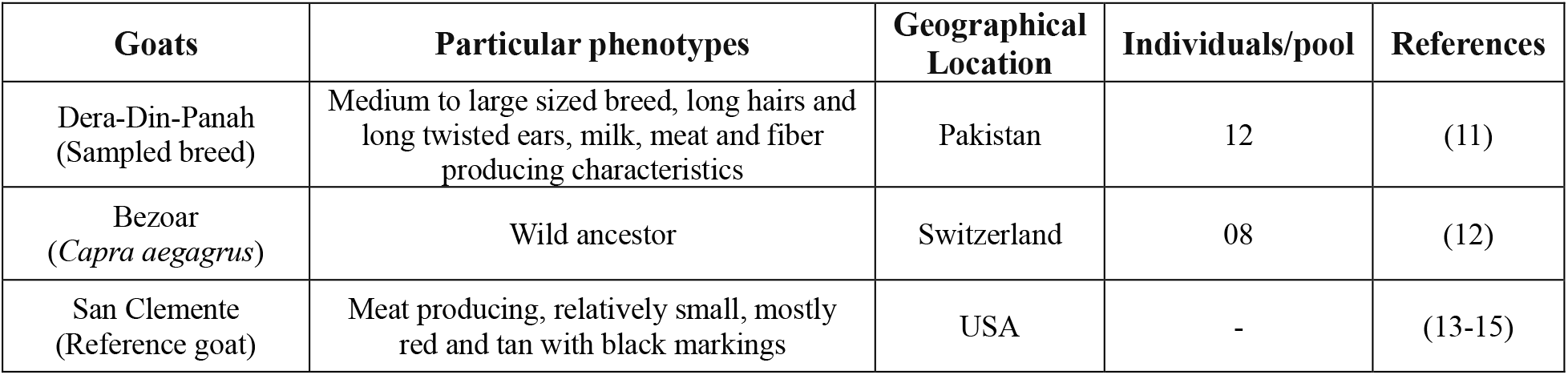
Particular features of sample DDP breed, wild ancestor (Bezoar) and reference goat (San Clemente)

| Goats | Particular phenotypes | Geographical Location | Individuals/pool | References |
| --- | --- | --- | --- | --- |
| Dera-Din-Panah (Sampled breed) | Medium to large sized breed, long hairs and long twisted ears, milk, meat and fiber producing characteristics | Pakistan | 12 | (11) |
| Bezoar (*Capra aegagrus*) | Wild ancestor | Switzerland | 08 | (12) |
| San Clemente (Reference goat) | Meat producing, relatively small, mostly red and tan with black markings | USA | - | (13-15) |

The pictures of sampled goat (DDP), reference goat (San Clemente) and wild ancestor (Bezoar) are shown (Fig. 1).

**Fig. 1.**
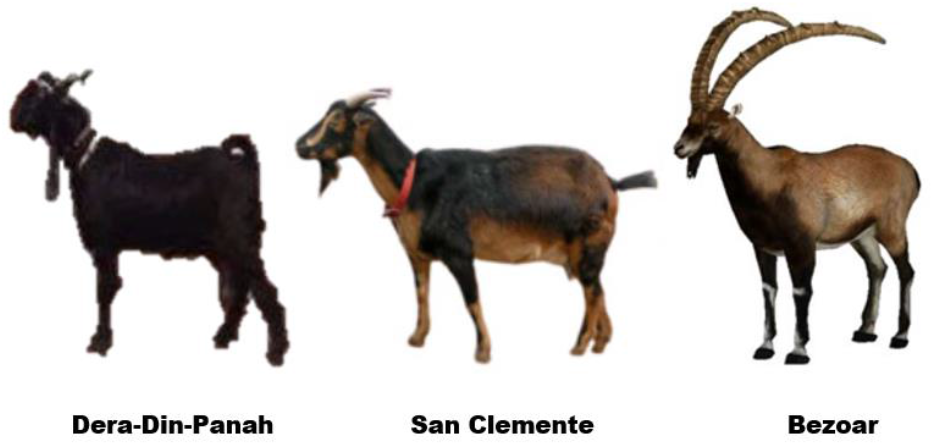
Representative individuals of the sampled breed (Dera-Din-Panah), reference (San clemente) and wild ancestor of the goat (Bezoar) are shown.

### Mapping and SNV calling of Pooled-Seq NGS data

Raw sequencing files (Fastq) of both goats (DDP, Bezoar) were quality filtered using Trimmomatic v 0.36 (16), mapped against goat reference genome using BWA-MEM algorithm, sorted on coordinate basis and duplicates in the samples were located and tagged using picard tool’s features, SortSam and MarkDuplicates respectively (17). First, SNPs, Indels and other complex variants along with the genotypes were called by applying Bayesian haplotype-based method implemented in FreeBayes v1.3.2 (18). Secondly, SNVs detection among DDP individuals was carried out by samtools mpileup (19). Popoolation2 v1.201 tool’s script, mpileup2sync.jar was utilized to synchronize the pileup file (20).

### Selective sweep detection using SFS statistical approaches

*SFS* based statistics except *TD* and *Pool – HMM* requires polarization of the allele i.e. the ancestral and derived state of the allele to be known which is achieved by the use of an outgroup species (21). Thus, the Bezoar goat was chosen as an outgroup and the VCF file was annotated with ancestral allele (AA) tag using BCFtools annotate function (19). This VCF was converted to haplotype file using data2haplohh function implemented in Relative Extended Haplotype Homozygosity (REHH) R-package (22) with parameter remove_multiple_markers = TRUE.

#### Tajima’s D, Fay & Wu’s H and Zeng’s E statistics

Using the calc_sfs_tests function of REHH R-package, *TD, H* and *E* statistics were computed which implies 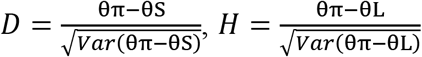, and 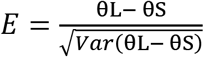 with sliding window of 150kb and an overlapping window size of 75kb (22, 23). Markers with more than 2 different alleles were discarded. Values were normalized by negative *Z*–transformation of each statistics 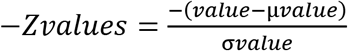.

#### Pool – HMM Statistics

A Hidden Markov Model (*HMM*) was applied on each autosome’s pileup file separately to detect allele frequencies and selective sweeps using a Python-based program, *Pool – HMM* (24), with parameters --min-coverage 3, --max-coverage 50, --min-quality 15 and number of haplotypes in the pool was set to 24. Data was visualized using a generic plot function of *R* along with the qplot function of ggplot2 package (25).

### Genomic selection pressure detection using reduced local-variability statistics

#### Pooled-Heterozygosity (*Hp*) Statistics

At each SNV position in sync pileup file, the major (nMAJ) and minor allele (nMIN) counts were observed using Popoolation2 v1.201 tool’s script, snp-frequency-diff.pl to further calculate *Hp* score that applies 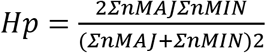 using an in-house Ruby script. The *Z*-transformation of each *Hp* value was performed, 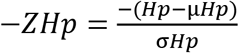 to normalize the data. Manhattan plots of all statistics were plotted using qqman package on R platform (26).

### Annotations of under-selection regions

The GCF_001704415.2 ARS1 reference goat genome assembly was utilized for annotations using Genome Data Viewer NCBI (27).

### Visualization of data using *R* software

R statistical packages e.g., ggplot2, qqman and plot function were used for the visualization of each statistics performed in the current analyses.

## Results

### Quality checks and mapping of WGS data

A total of 85,703,935,001 base counts were sequenced with 30x genome coverage. After trimming and filtering, 618,236,192 clean reads were mapped to the ARS1 reference genome assembly. Overall, the GC content (42%), sequence length (30-151bp), no overrepresentation and no sequence tagged as poor quality illustrates that high quality sequence was obtained in this study (Fig. S1).

### Whole-genome selection signature scan and SNPs detection

Variants in VCF were supplemented with ancestral allele information that resulted in 31,273,657 markers out of which 30,745,656 markers harboring 3,289 windows were obtained after applying SFS statistics. Based on the computation of *TD* statistics (–*ZTD* ≥ 4.7, μ=0.598 and σ=0.341), 59 windows were identified having 21,760 significant SNPs. For *H* statistics (–*ZH* ≥ 6, μ=-0.104 and σ=0.659), 67 under-selection windows were detected that spanned 26,117 selection markers while *E* statistics –*ZE* ≥ 2.5, μ=0.694 and *σ*=0.791 resulted in 57 candidate windows harboring 53,535 number of significant SNP markers. *Pool – HMM* with threshold ≥ 12, μ=7.338 and *σ*=3.520 yielded 76 selection windows. Similarly, the *Hp* analysis (–*ZHp* ≥ 5, μ=0.321 and *σ*=0.028) called 32,581,969 SNPs residing on 32,839 windows of which 80 windows and 30,045 markers were found under selection pressure. The above-mentioned significant threshold values for each statistical measure were assigned on the basis of published data as well as on the basis of in-house adopted stringent cutoff values for better selection signature hits. The SNPs frequency distribution graph of each statistics across all the 29-autosomes of DDP breed is shown in Fig. 2

**Fig. 2.**
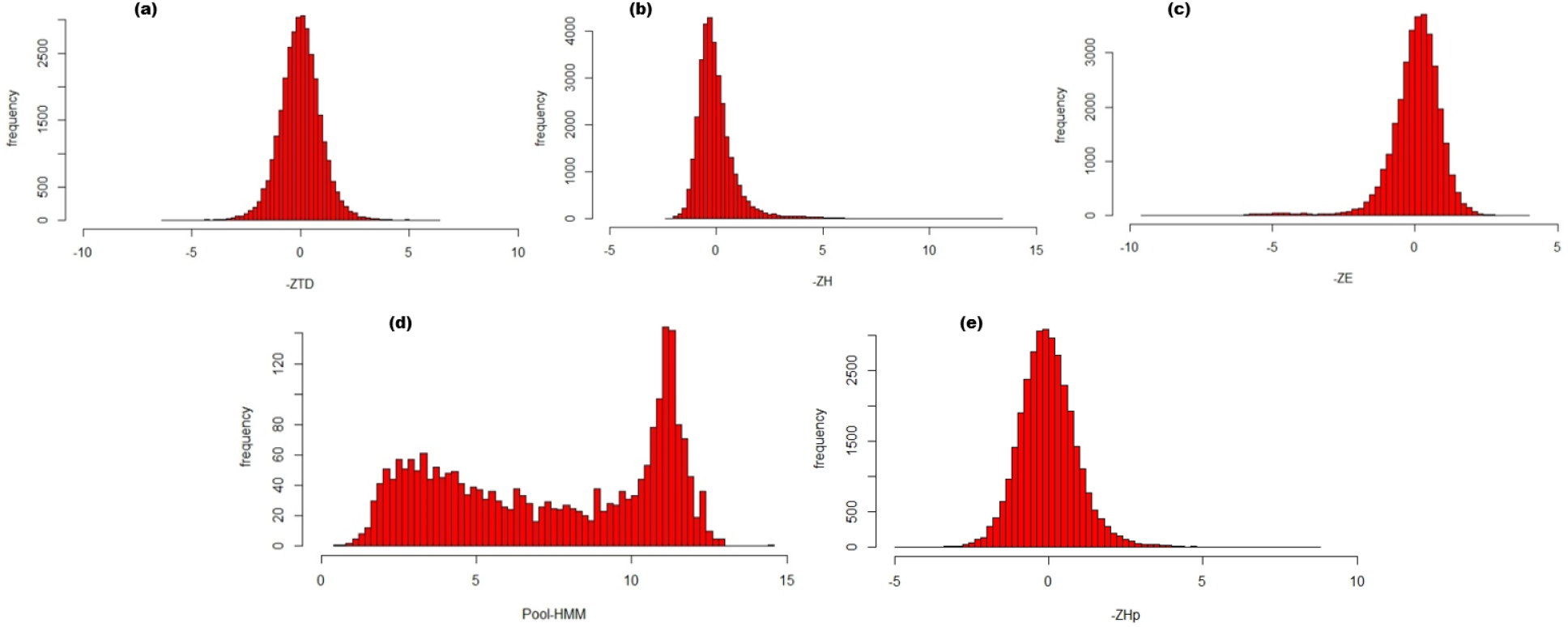
Distribution of SNPs against –*ZTD, –ZH, –ZE, Pool – HMM, and – ZHp* (a, b, c, d, e respectively) signals across all autosomes of DDP goat

### Selection windows detected by *TD, H* and *E* statistics

The *TD* analysis detected 59 windows with slightly negative values indicating intermediate signals spanning 63 genes, 21 LOC symbols and 11 regions that are devoid of genes. Variation in allele frequency before fixation was also observed by *H*-statistics of which the negative value signifies an excess of derived polymorphism with respect to neutral expectations. Out of sixty-seven under selection regions, 22 windows do not harbor any genes while rest of comprises 60 genes, and 32 LOC symbols. Similarly, the *E*-statistics identified 109 genes, 57 LOC symbols, and 3 regions have no genes. The complete list of selection signature windows identified from three *SFS* methods is provided (Table S1). Manhattan plots are displayed in Fig. 3 showing the distribution of SNPs across the whole genome against the –*ZTD*, -*ZH*, and –*ZE* scores.

**Fig. 3.**
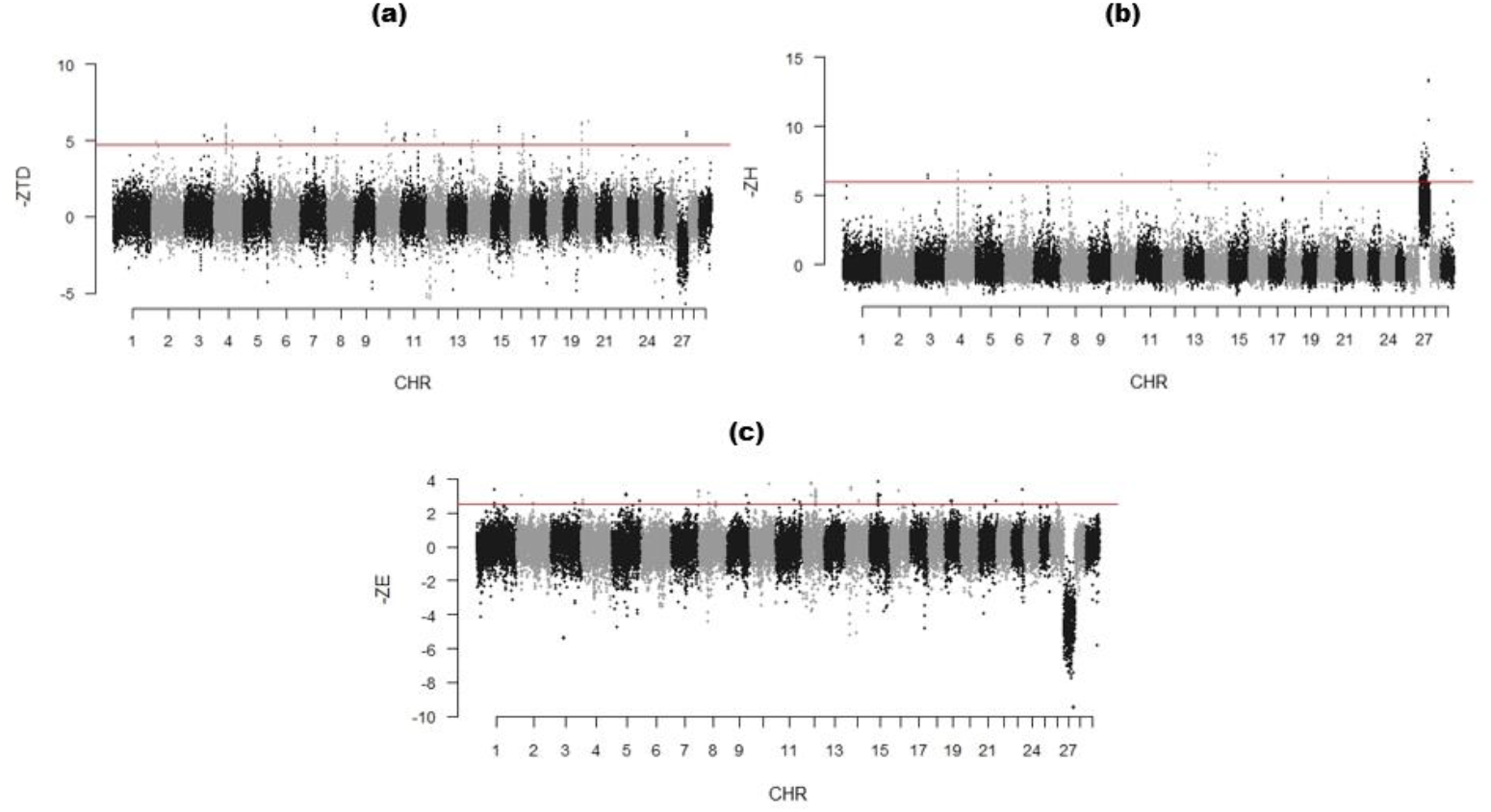
Manhattan plots of -ZTD, -ZH, and -ZE values across whole genome with window size 150kb and step size of 75kb. Red bar indicates the significant threshold of –*ZTD* ≥ 4.7, –*ZH* ≥ 6, and -*ZE* ≥ 2.5.

### Genomic selection pressure regions detected by *Pool — HMM*

Seventy-six top selection windows above the threshold line in (Fig. 4) were identified using *Pool – HMM*. The .stat file reporting the maximum of posterior probabilities of hidden state along the windows is provided (Table S2). The top hits harbor 163 genes and 73 LOC symbols while 19 regions do not contain any genes.

**Fig. 4.**
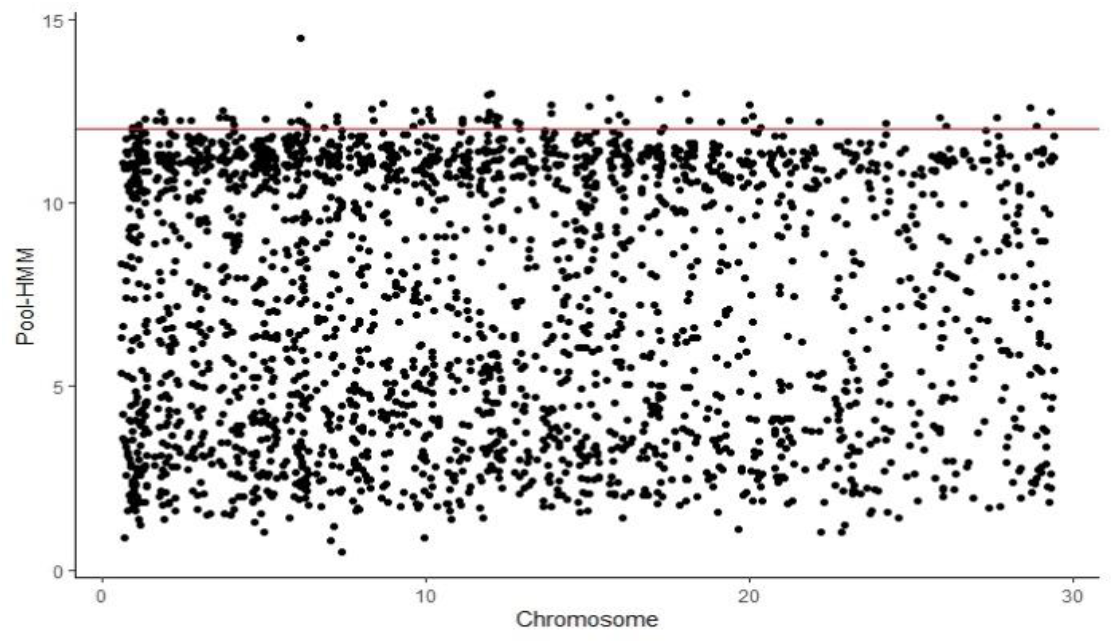
Scatter-plot of *Pool – HMM* values across the whole genome. Red bar signifies the threshold of *Pool – HMM* ≥ 12

Secondly, allele counts and their probabilities against each autosome of DDP goat were also plotted individually in (Fig. S2). It represents the posterior probability of loci under selection (hidden state) that is calculated for each genomic locus with the help of forward–backward algorithm implemented in *Pool – HMM*, which generated the .post suffix resultant files for better understanding of the regions under selection pressure in each chromosome.

### Genomic scan for positive sweeps using *Hp* statistics

*Hp* analysis was computed on major and minor alleles of significant SNPs that resulted in 80 under-selection windows with –*ZHp ≥ 5*. Out of these windows, 13 regions do not contain any genes while remaining regions composed of 122 genes, and 41 LOC symbols. Complete genomic selection signature list is provided (Table S3) while Fig. 5 displays genome wide distribution of selection signals plotted along DDP goat autosomes.

**Fig. 5.**
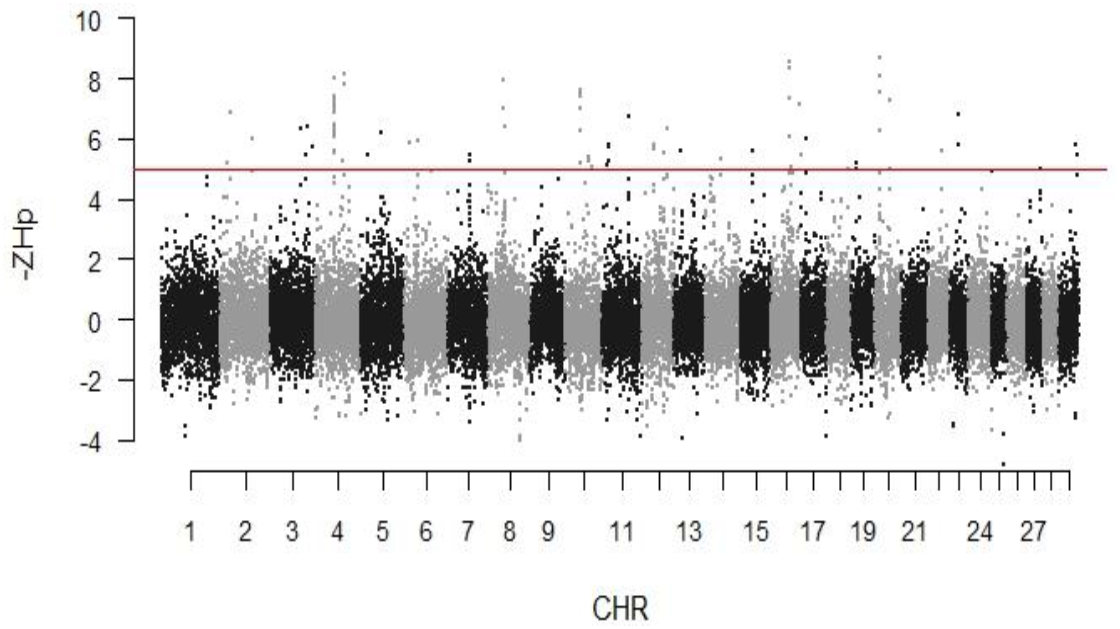
Manhattan plot of Z-transformed values of *Hp* with window size 150kb and 75kb step size. The horizontal bar displays cutoff value –*ZHp ≥ 5* used for extracting outliers.

### Annotation of selection signals

Highly significant regions were further annotated with genes that are majorly associated with body height, weight, milk production, fecundity rate/reproduction, skeleton development, and immunity. A few genes also showed association with pigmentation, olfaction, wool production, and adaptation capability to various environments (Table 2).

**Table 2.**
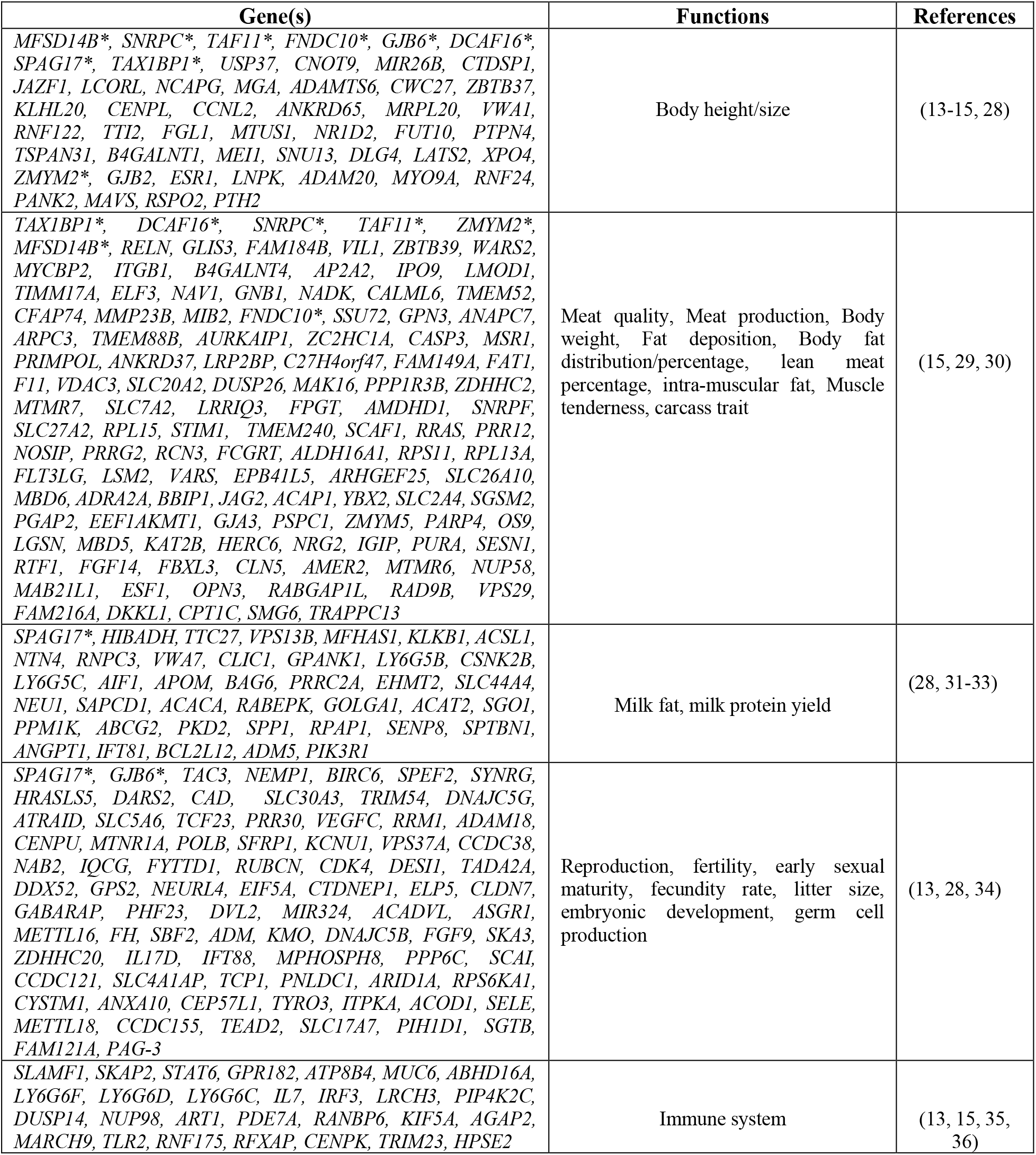
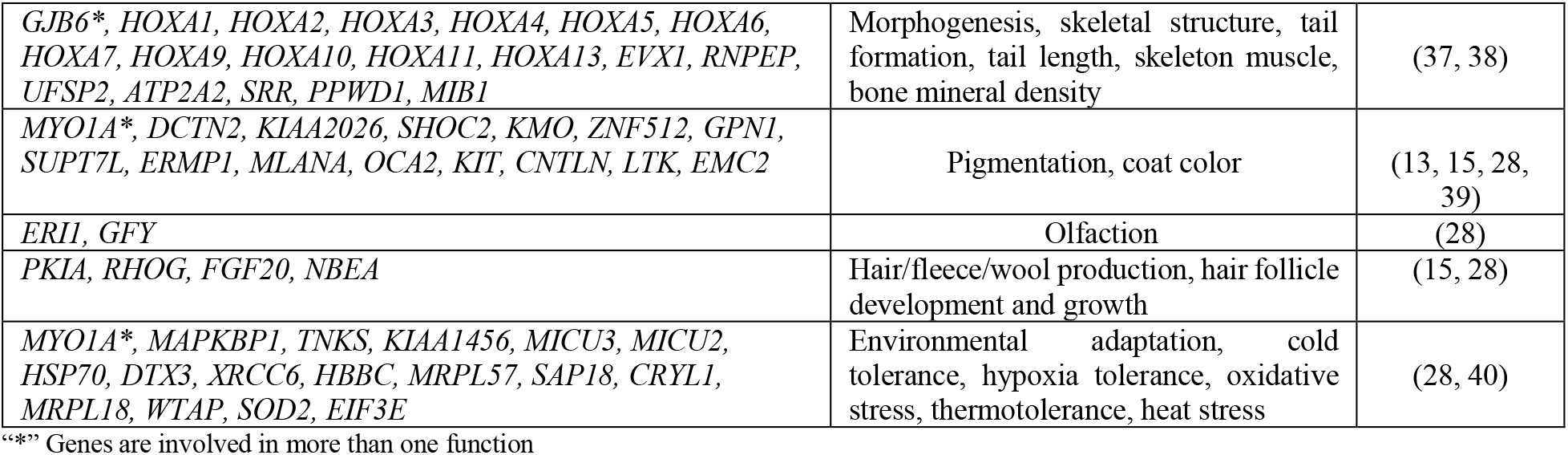
List of under-selection candidate genes from all five statistical methods and their association with important traits in DDP goat.

### Common putative selection windows

Fine mapping of under-selection regions was further carried out for extracting the putative windows that were captured by more than one aforementioned applied statistical approaches to get more accurate and reliable windows under genomic selection pressure in this goat breed. Thus, thirty-three common sweeps were considered as putative regions that covered 110 genes and 38 LOCs (Table 3) which are significantly associated with the vested traits (Table 1) of this particular goat breed.

**Table 3.**
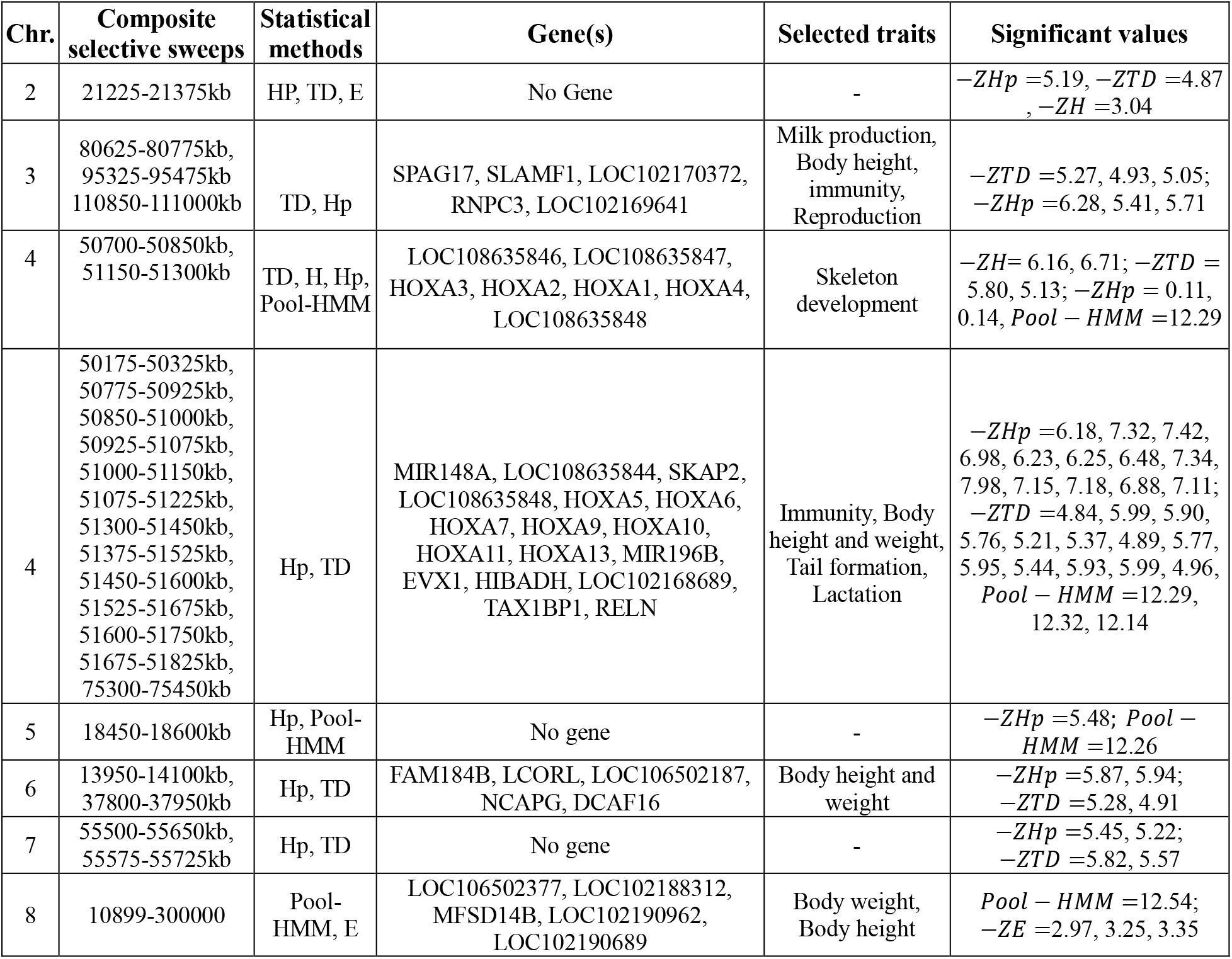
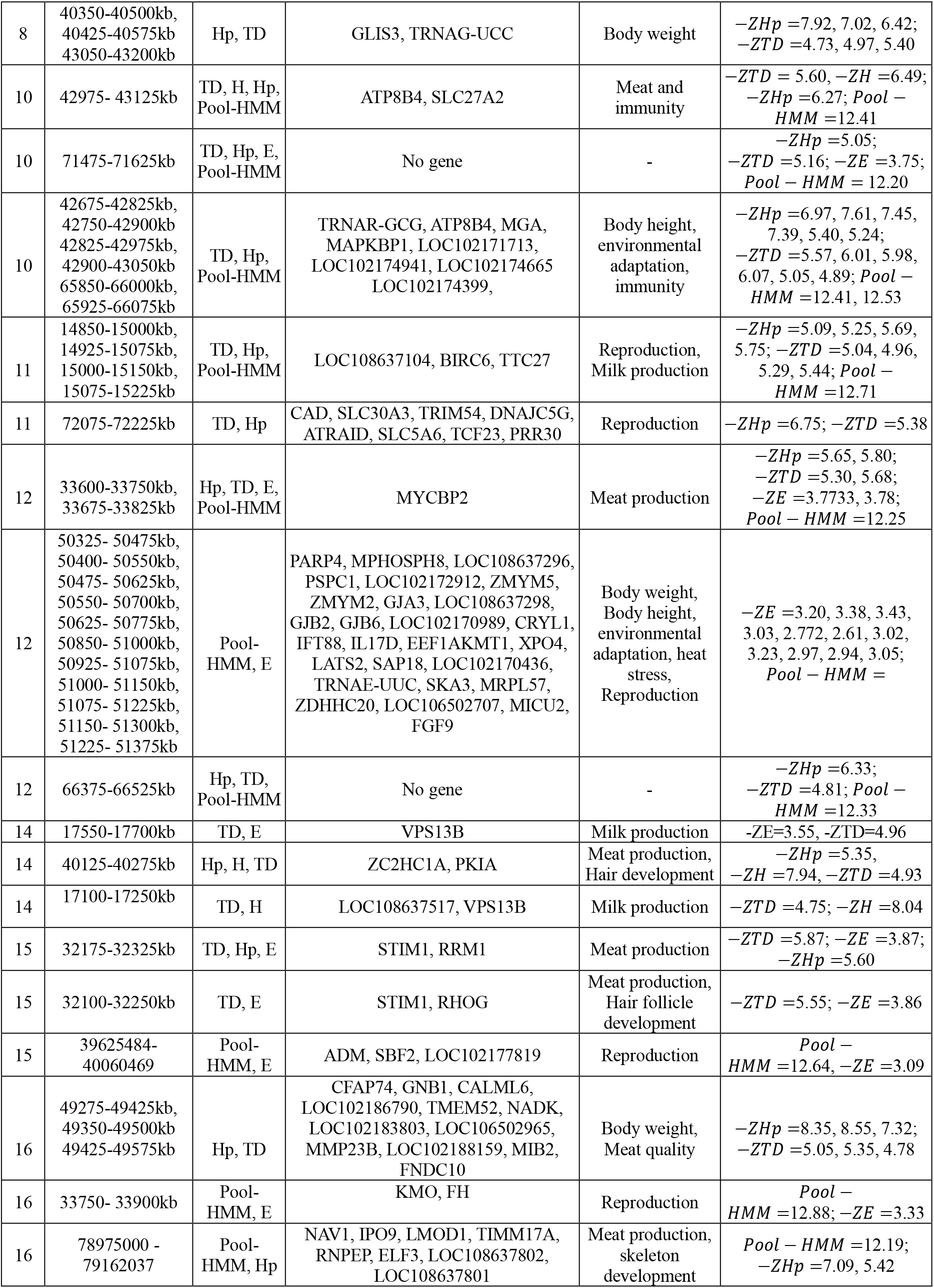
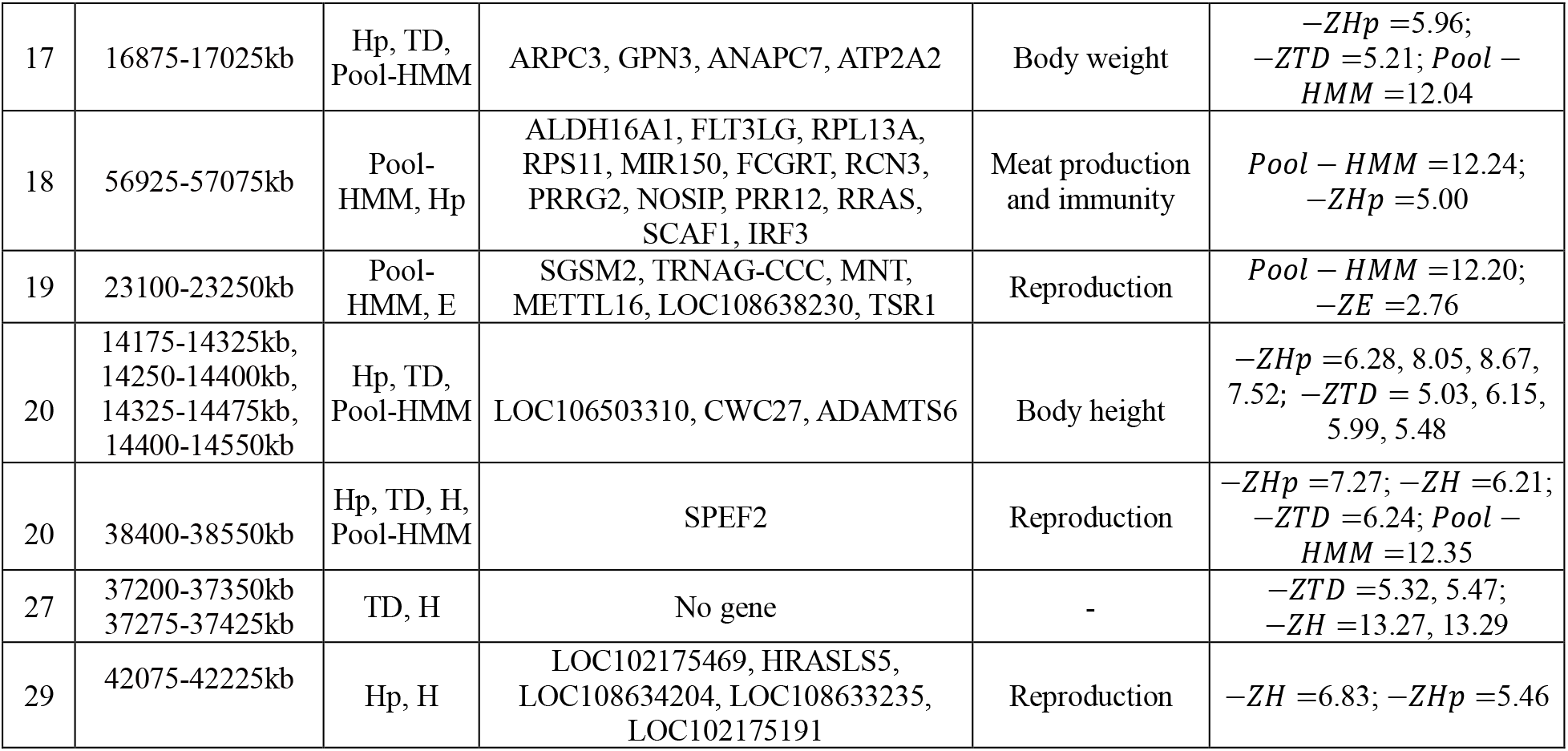
Top captured selection signals identified via either *TD, H, E, Pool – HMM* or *Hp* statistics.

## Discussion

Detection of genetic loci affecting valuable economic traits in livestock has long been of the interest to breeders and goat farmers for controlling and improving their characteristics. The current study was designed to detect the selection signatures in whole-genome pool-seq data of DDP goat breed by applying *TD,H,E,Pool – HMM*, and *Hp* statistics. DDP goat is famous for meat, milk, and wool production. We extracted 41 common genes that are functionally related to body weight/meat quality, for instance, *STIM1*, and *RRM1* genes were previously reported to be under-selection in Swiss goat breeds using Runs of Homozygosity (*ROH*) approach (8). Similarly, *IPO9, TIMM17A*, has been under-selection in Nellore cattle with significant *p*-values of 0.01820, and 0.01768 respectively. *MIB2* which is under-selection in our DDP breed is a muscle E3 ubiquitin ligase containing an active E3 RING-finger domain that is required for myoblast fusion in embryos (41).

The genes (*RELN, NAV1, LMOD1, ELF3*) are associated with body weight and body mass index in humans (28). Out of total 40 milk production genes, only 4 genes (*VPS13B, TTC27, RNPC3*, and *SPAG17)* harbor the common putative windows. Four SNPs at 15~15.3 Mb region within *TTC27* gene were significant (FDR < 5%) in Bovine genome (42), while *TTC27* along with *VPS13B* gene was found to be under-selection in Pakistani Teddy goat breed (13). Only two of five fiber production genes (*PKIA* and *RHOG*) captured by more than one statistical method used in the current research. *PKIA* gene with –*ZHp = 7.520* was under high selection in Teddy goat (13) while currently in DDP goat it is significantly associated with score –*ZHp = 5.35* (Table S3). Interestingly, *RHOG* was found as candidate gene involved in number of hair whorls on right side in horses thus evidencing the curly hair in DDP goat as well (43).

In short, outlier loci and genes identified in our study showed the high degree of genetic diversity within DDP goat breed, but still further functional studies are required to depict the underlying mechanism of candidate genes and to utilize this valuable information for future quantitative trait loci mapping for the goat breeding.

## Acknowledgements

The authors are thankful to all goat owners who donated the samples of their animals. Also, we are grateful to Prof. Dr. Tosso Leeb, Vidhya Jagannathan and Jan Henkel for their consistent support in accomplishing this research endeavor. We are obliged to University of Bern, Switzerland for providing us with the NGS platform for conducting this high-throughput sequencing experiments.

## References

1. Li J, Zhang Y. Advances in research of the origin and domestication of domestic animals. Biodiversity Science. 2009;17(4):319.

2. Bertolini F, Servin B, Talenti A, Rochat E, Kim ES, Oget C, et al. Signatures of selection and environmental adaptation across the goat genome post-domestication. Genetics Selection Evolution. 2018;50(1):1–24.

3. Guo J, Tao H, Li P, Li L, Zhong T, Wang L, et al. Whole-genome sequencing reveals selection signatures associated with important traits in six goat breeds. Scientific reports. 2018;8(1):1–11.

4. Gurgul A, Jasielczuk I, Semik-Gurgul E, Pawlina-Tyszko K, Stefaniuk-Szmukier M, Szmatoła T, et al. A genome-wide scan for diversifying selection signatures in selected horse breeds. PLoS One. 2019;14(1):e0210751.

5. Kim J-Y, Jeong S, Kim KH, Lim W-J, Lee H-Y, Kim N. Discovery of genomic characteristics and selection signatures in Korean indigenous goats through comparison of 10 goat breeds. Frontiers in genetics. 2019;10:699.

6. Muner R, Bilal G, Moaeen-ud-Din M, Reecy J, Khan M. 36 Morphometric measurements and body weight is determined by breed, age and sex among Punjab goat breeds of Pakistan. Journal of Animal Science. 2018;96(Suppl 3):453.

7. Ronen R, Udpa N, Halperin E, Bafna V. Learning natural selection from the site frequency spectrum. Genetics. 2013;195(1):181–93.

8. Signer-Hasler H, Henkel J, Bangerter E, Bulut Z, Drögemüller C, Leeb T, et al. Runs of homozygosity in Swiss goats reveal genetic changes associated with domestication and modern selection. Genetics Selection Evolution. 2022;54(1):1–11.

9. Anand S, Mangano E, Barizzone N, Bordoni R, Sorosina M, Clarelli F, et al. Next generation sequencing of pooled samples: guideline for variants’ filtering. Scientific reports. 2016;6(1):1–9.

10. Lynch M, Bost D, Wilson S, Maruki T, Harrison S. Population-genetic inference from pooled-sequencing data. Genome biology and evolution. 2014;6(5):1210–8.

11. Afzal M, Naqvi A. Livestock resources of Pakistan: present status and future trends. Quarterly Science Vision. 2004;9(1):1–2.

12. Takada T, Kikkawa Y, Yonekawa H, Kawakami S, Amano T. Bezoar (Capra aegagrus) is a matriarchal candidate for ancestor of domestic goat (Capra hircus): evidence from the mitochondrial DNA diversity. Biochemical Genetics. 1997;35(9):315–26.

13. Saif R, Henkel J, Mahmood T, Ejaz A, Ahmad F, Zia S. Detection of whole genome selection signatures of Pakistani Teddy goat. Molecular Biology Reports. 2021;48(11):7273–80.

14. Saif R, Henkel J, Jagannathan V, Drögemüller C, Flury C, Leeb T. The LCORL locus is under selection in large-sized Pakistani goat breeds. Genes. 2020;11(2):168.

15. Saif R, Mahmood T, Ejaz A, Fazlani SA, Zia S. Whole-genome selective sweeps analysis in Pakistani Kamori goat. Gene Reports. 2022;26:101429.

16. Bolger AM, Lohse M, Usadel B. Trimmomatic: a flexible trimmer for Illumina sequence data. Bioinformatics. 2014;30(15):2114–20.

17. Li H, Durbin R. Fast and accurate long-read alignment with Burrows–Wheeler transform. Bioinformatics. 2010;26(5):589–95.

18. Garrison E, Marth G. Haplotype-based variant detection from short-read sequencing. arXiv preprint arXiv:12073907. 2012.

19. Danecek P, Bonfield JK, Liddle J, Marshall J, Ohan V, Pollard MO, et al. Twelve years of SAMtools and BCFtools. Gigascience. 2021;10(2):giab008.

20. Popoolation2 [Available from: https://sourceforge.net/projects/popoolation2/.

21. Klassmann A, Gautier M. Detecting selection using extended haplotype homozygosity (EHH)-based statistics in unphased or unpolarized data. PloS one. 2022;17(1):e0262024.

22. Gautier M, Vitalis R. rehh: an R package to detect footprints of selection in genome-wide SNP data from haplotype structure. Bioinformatics. 2012;28(8):1176–7.

23. Zeng K, Fu Y-X, Shi S, Wu C-I. Statistical tests for detecting positive selection by utilizing high-frequency variants. Genetics. 2006;174(3):1431–9.

24. Boitard S, Kofler R, Françoise P, Robelin D, Schlötterer C, Futschik A. Pool-hmm: A Python program for estimating the allele frequency spectrum and detecting selective sweeps from next generation sequencing of pooled samples. Molecular ecology resources. 2013;13(2):337–40.

25. Tollefson M. Graphics with the ggplot2 Package: An Introduction. Visualizing Data in R 4: Springer; 2021. p. 281–93.

26. manhattan: Creates a manhattan plot [Available from: https://rdrr.io/cran/qqman/man/manhattan.html.

27. NCBI: Genome Data Viewer [Available from: https://www.ncbi.nlm.nih.gov/genome/gdv?org=capra-hircus&group=bovidae.

28. GeneCards®: The Human Gene Database [Available from: https://www.genecards.org/.

29. Verardo LL, Sevón-Aimonen M-L, Serenius T, Hietakangas V, Uimari P. Whole-genome association analysis of pork meat pH revealed three significant regions and several potential genes in Finnish Yorkshire pigs. BMC genetics. 2017;18(1):1–15.

30. Zhang G, Fan Q, Wang J, Zhang T, Xue Q, Shi H. Genome-wide association study on reproductive traits in Jinghai Yellow Chicken. Animal reproduction science. 2015;163:30–4.

31. Buaban S, Lengnudum K, Boonkum W, Phakdeedindan P. Genome-wide association study on milk production and somatic cell score for Thai dairy cattle using weighted single-step approach with random regression test-day model. Journal of Dairy Science. 2022;105(1):468–94.

32. Dai W-t, Zou Y-x, White RR, Liu J-x, Liu H-y. Transcriptomic profiles of the bovine mammary gland during lactation and the dry period. Functional & integrative genomics. 2018;18(2):125–40.

33. Seo M, Lee H-J, Kim K, Caetano-Anolles K, Jeong JY, Park S, et al. Characterizing milk production related genes in Holstein using RNA-seq. Asian-Australasian journal of animal sciences. 2016;29(3):343.

34. Melo TPd, De Camargo GMF, De Albuquerque LG, Carvalheiro R. Genome-wide association study provides strong evidence of genes affecting the reproductive performance of Nellore beef cows. PLoS One. 2017;12(5):e0178551.

35. Fontanesi L, Galimberti G, Calò D, Fronza R, Martelli P, Scotti E, et al. Identification and association analysis of several hundred single nucleotide polymorphisms within candidate genes for back fat thickness in Italian Large White pigs using a selective genotyping approach. Journal of Animal Science. 2012;90(8):2450–64.

36. Xia X, Zhang S, Zhang H, Zhang Z, Chen N, Li Z, et al. Assessing genomic diversity and signatures of selection in Jiaxian Red cattle using whole-genome sequencing data. BMC genomics. 2021;22(1):1–11.

37. Fan B, Onteru SK, Du Z-Q, Garrick DJ, Stalder KJ, Rothschild MF. Genome-wide association study identifies loci for body composition and structural soundness traits in pigs. PloS one. 2011;6(2):e14726.

38. Yuan Z, Liu E, Liu Z, Kijas J, Zhu C, Hu S, et al. Selection signature analysis reveals genes associated with tail type in Chinese indigenous sheep. Animal genetics. 2017;48(1):55–66.

39. Henkel J, Saif R, Jagannathan V, Schmocker C, Zeindler F, Bangerter E, et al. Selection signatures in goats reveal copy number variants underlying breed-defining coat color phenotypes. PLoS genetics. 2019;15(12):e1008536.

40. Xu K, Niu Q, Zhao H, Du Y, Jiang Y. Transcriptomic analysis to uncover genes affecting cold resistance in the Chinese honey bee (Apis cerana cerana). PLoS One. 2017;12(6):e0179922.

41. Ribot C, Soler C, Chartier A, Al Hayek S, Naït-Saïdi R, Barbezier N, et al. Activation of the ubiquitin-proteasome system contributes to oculopharyngeal muscular dystrophy through muscle atrophy. PLoS genetics. 2022;18(1):e1010015.

42. Nayeri S, Schenkel F, Fleming A, Kroezen V, Sargolzaei M, Baes C, et al. Genome-wide association analysis for β-hydroxybutyrate concentration in Milk in Holstein dairy cattle. BMC genetics. 2019;20(1):1–17.

43. Lima DFPdA, da Cruz VAR, Pereira GL, Curi RA, Costa RB, de Camargo GMF. Genomic Regions Associated with the Position and Number of Hair Whorls in Horses. Animals. 2021;11(10):2925.

