## Supplementary Figures for "Genomic Selection Pressure Discovery using Site-Frequency Spectrum & Reduced Local-Variability Statistics in Pakistani Dera-Din-Panah Goat"

**Supplementary figure 1**

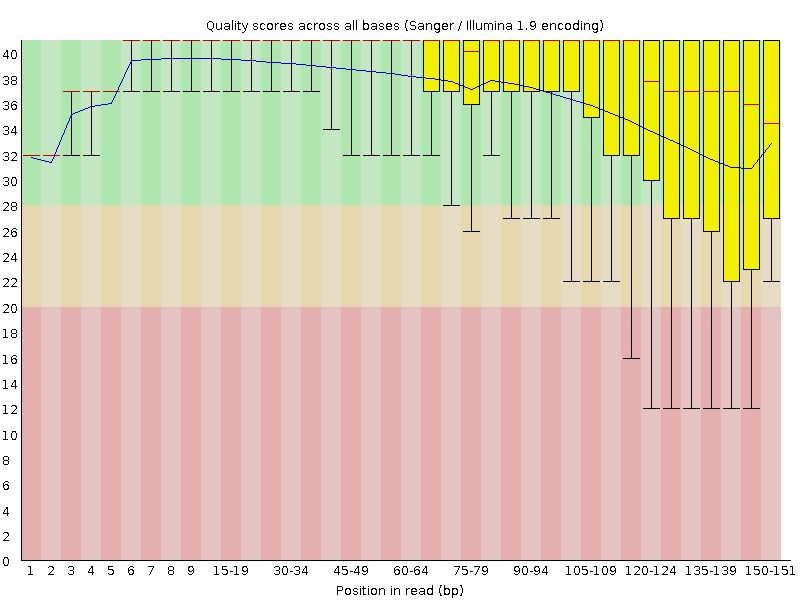

**Fig.S1** Position on x-axis vs. phred score on y-axis was plotted as per base sequence quality graph by FastQC software.

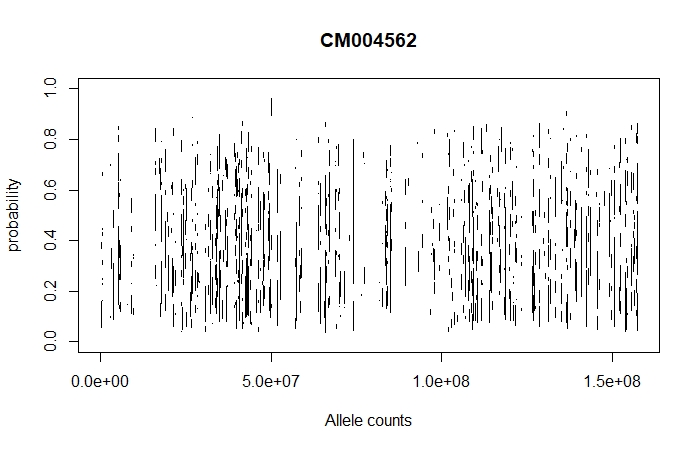

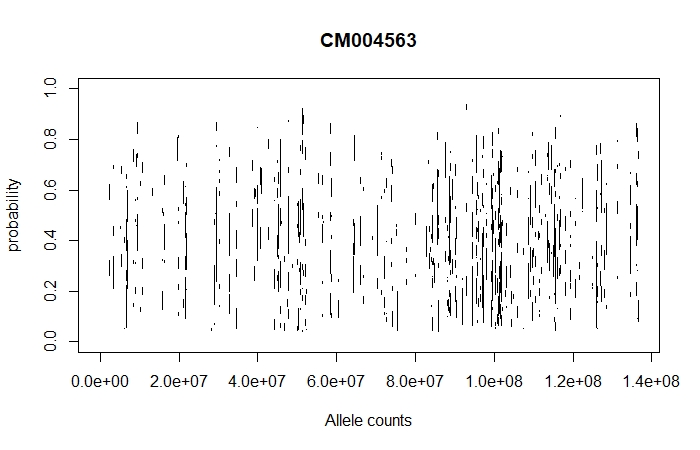

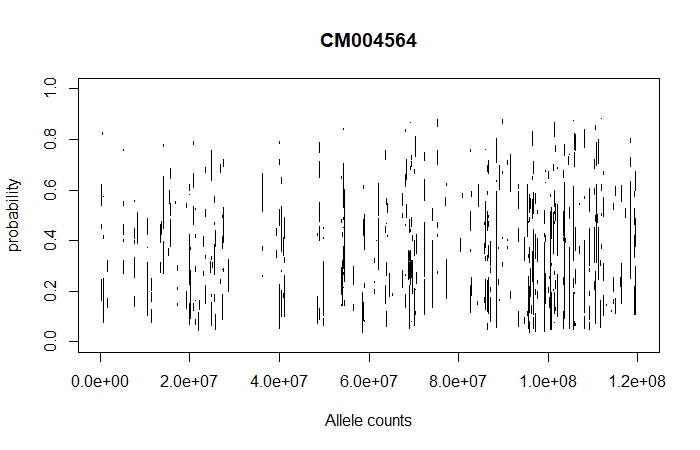

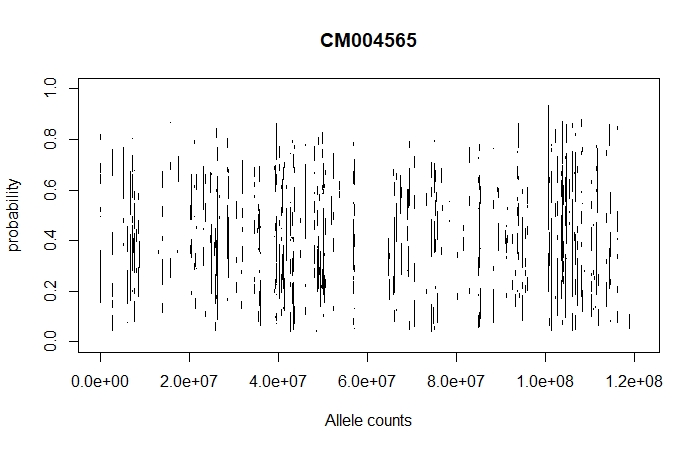

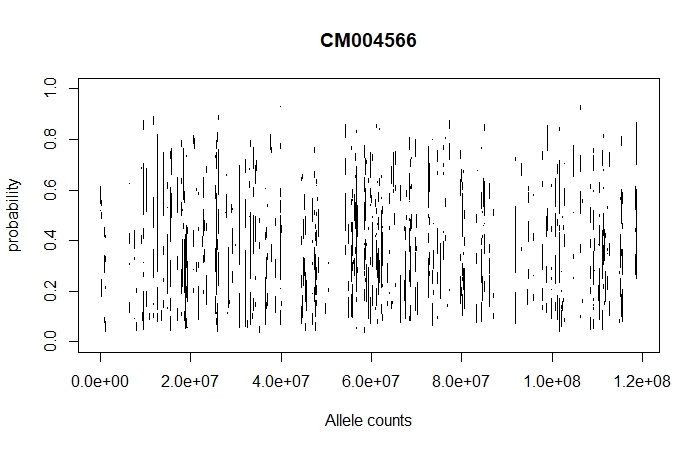

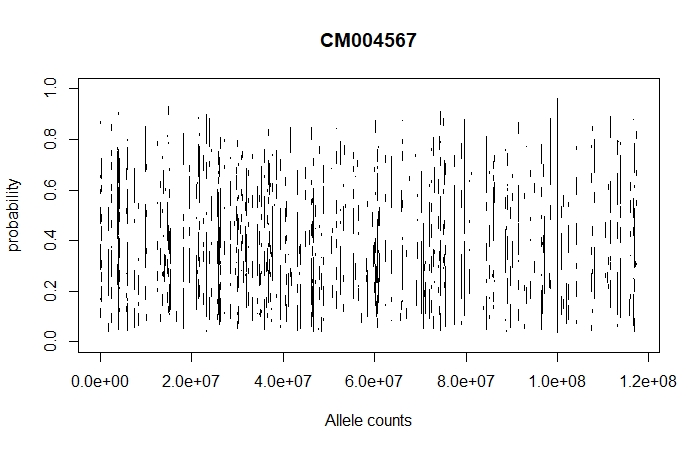

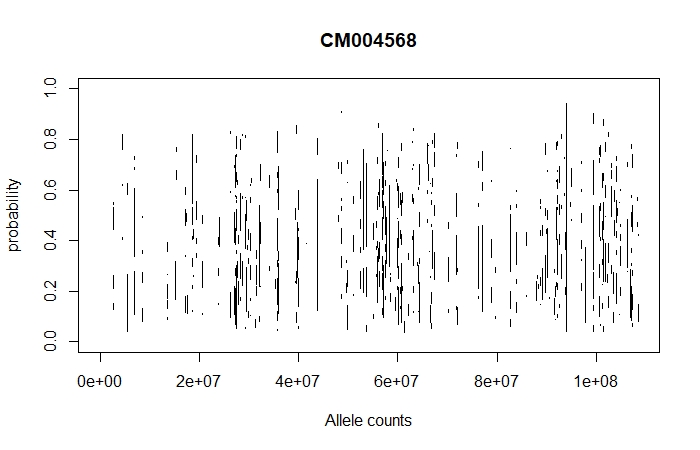

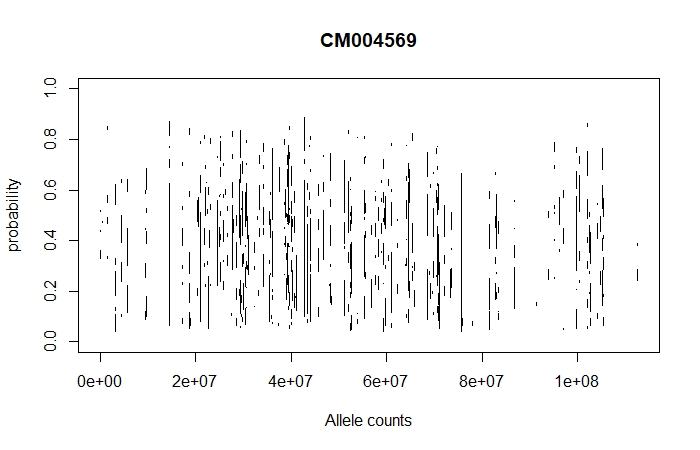

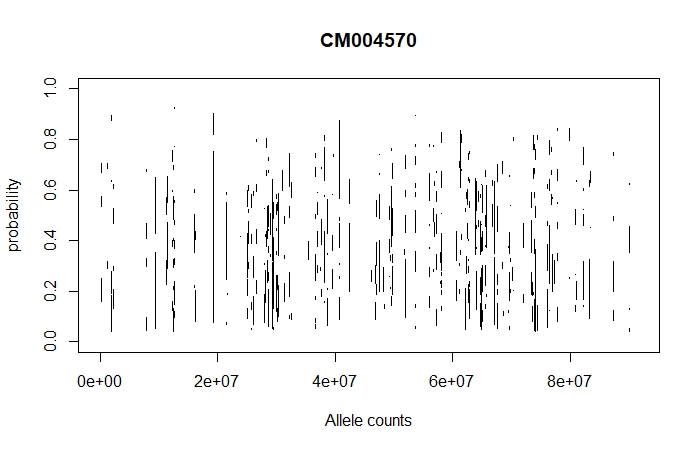

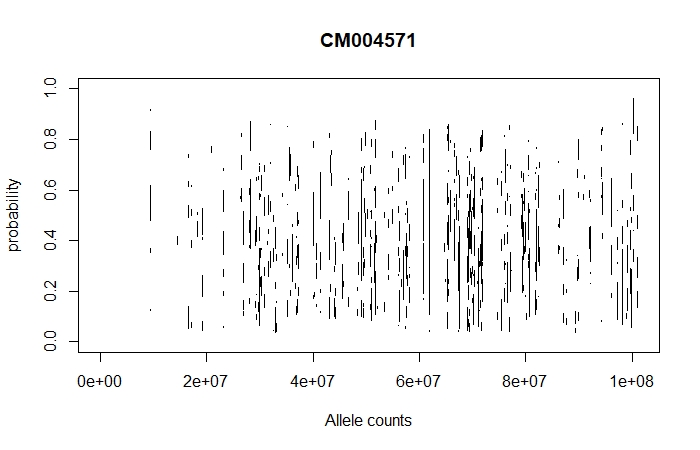

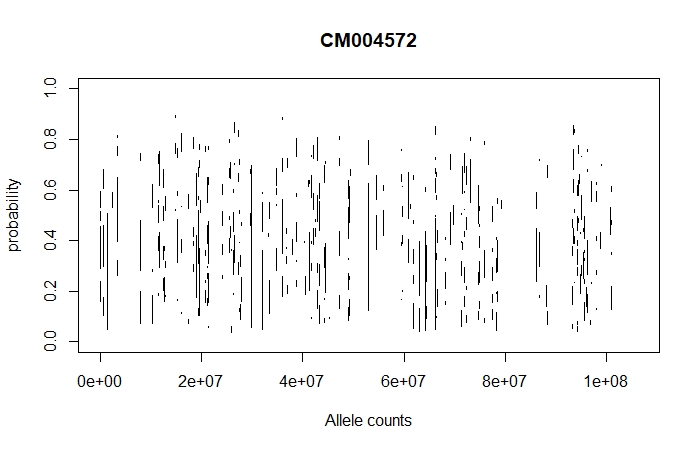

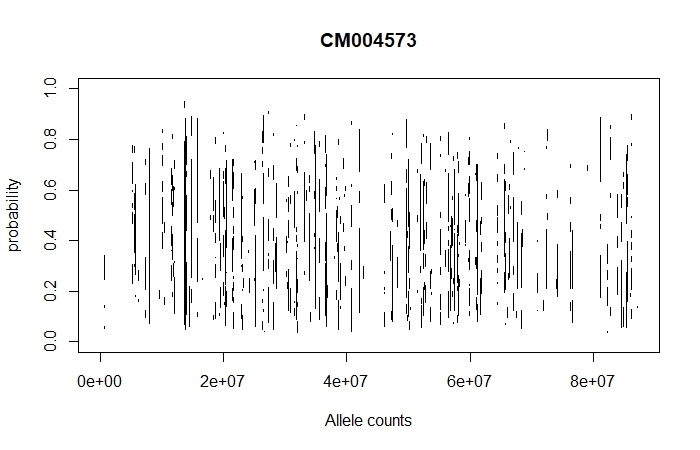

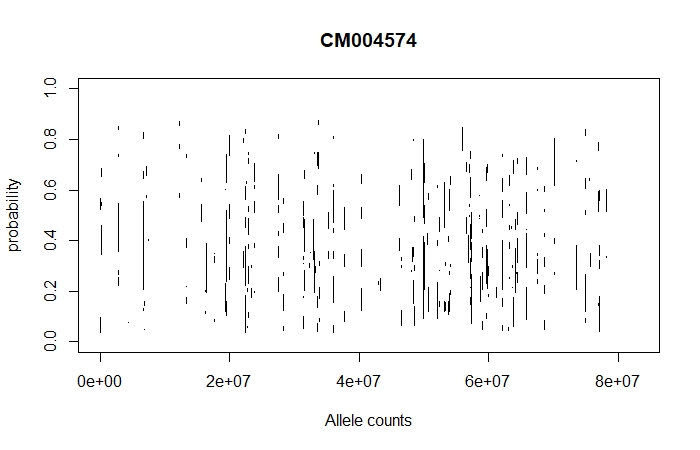

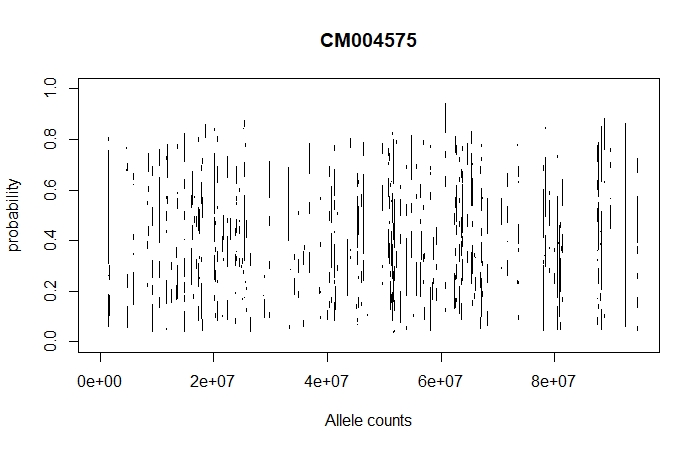

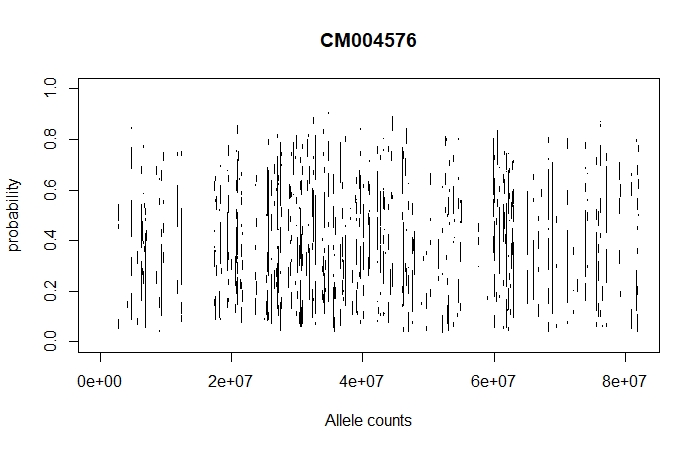

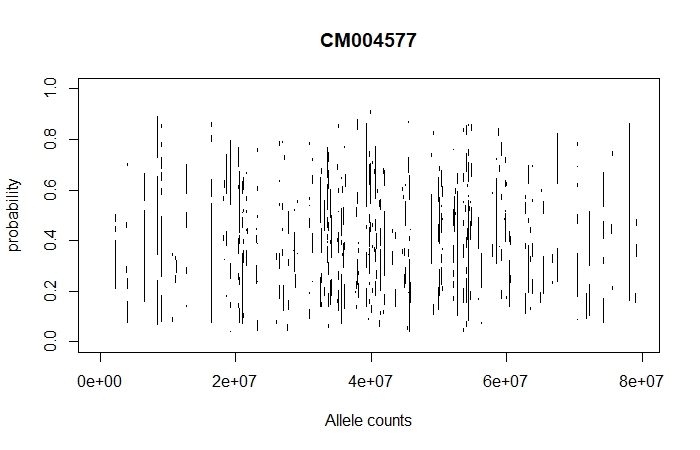

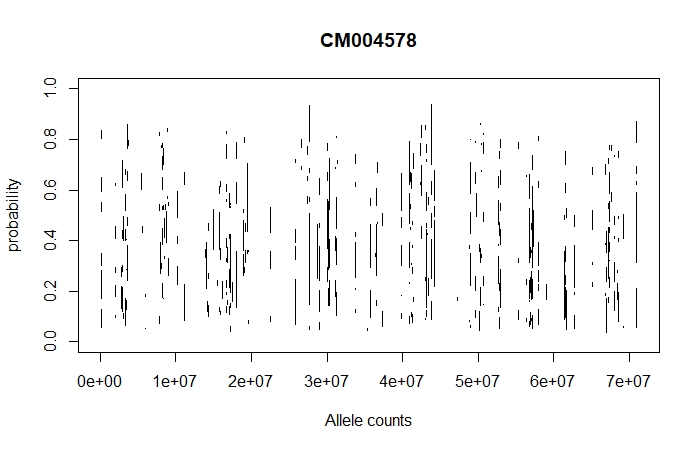

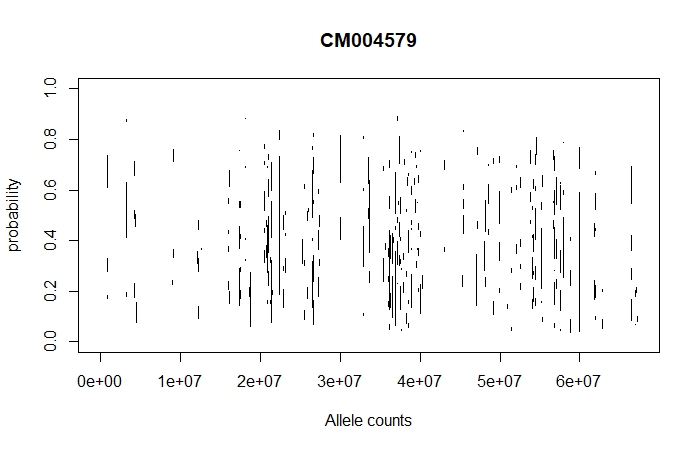

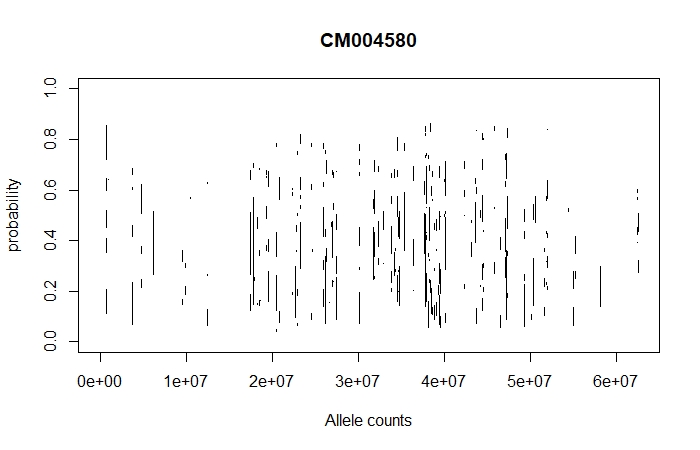

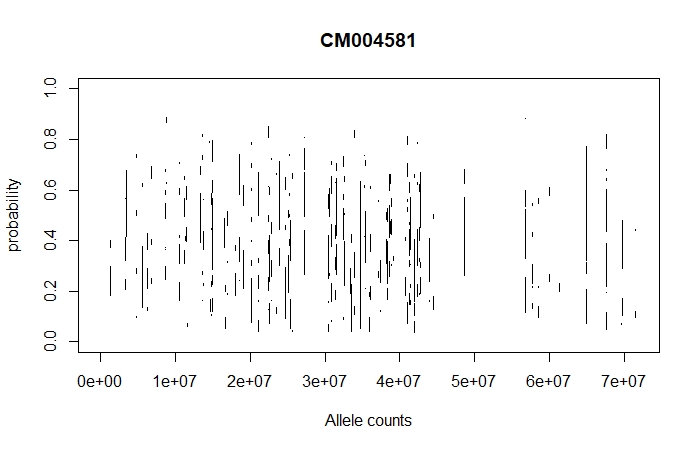

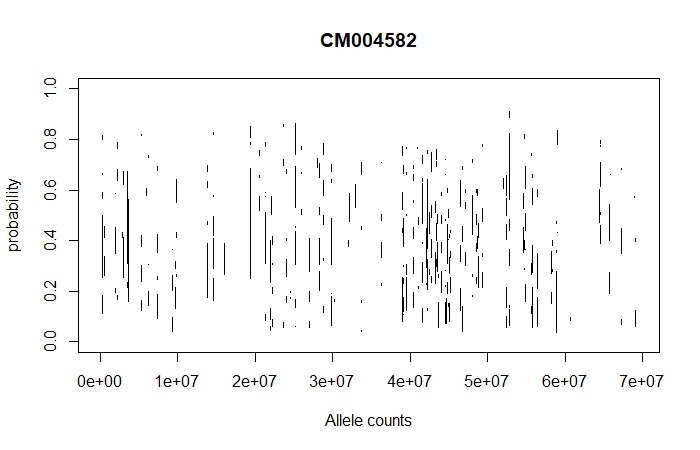

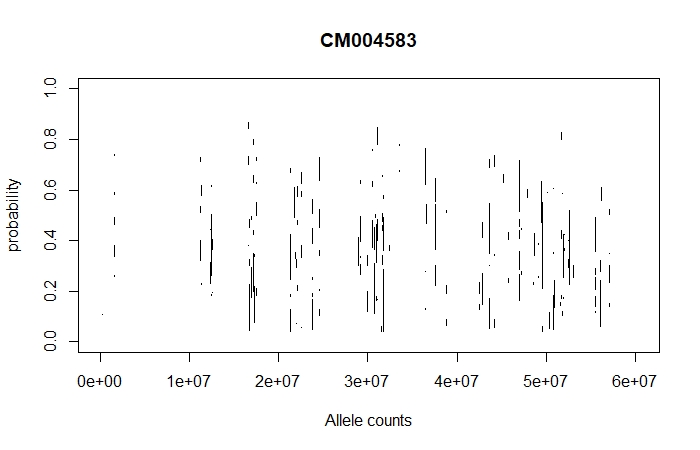

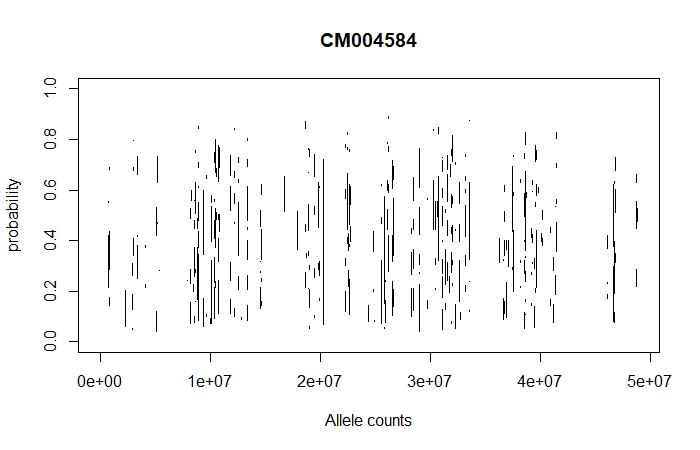

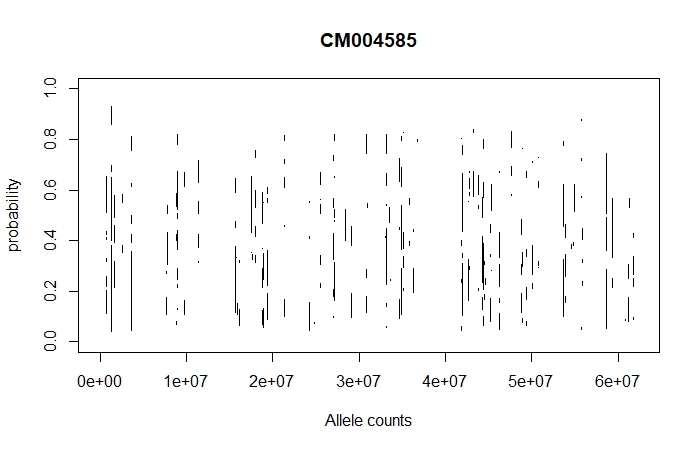

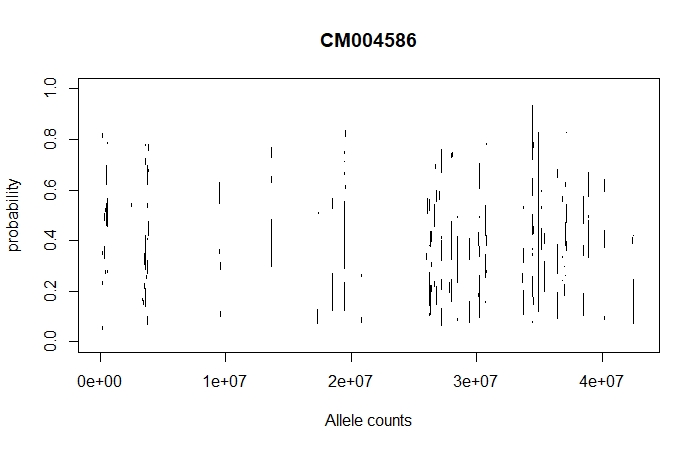

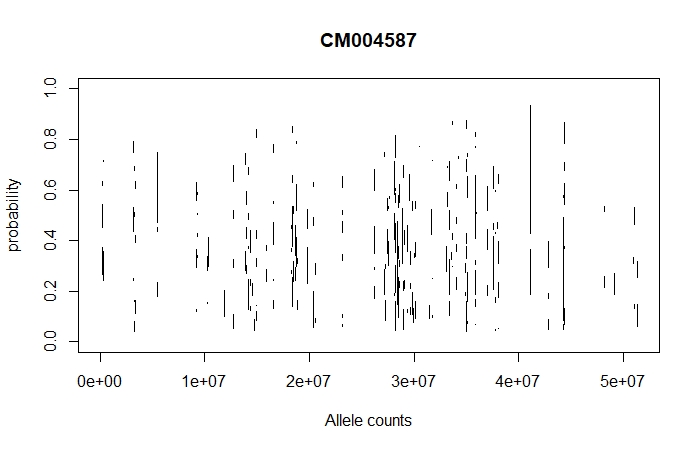

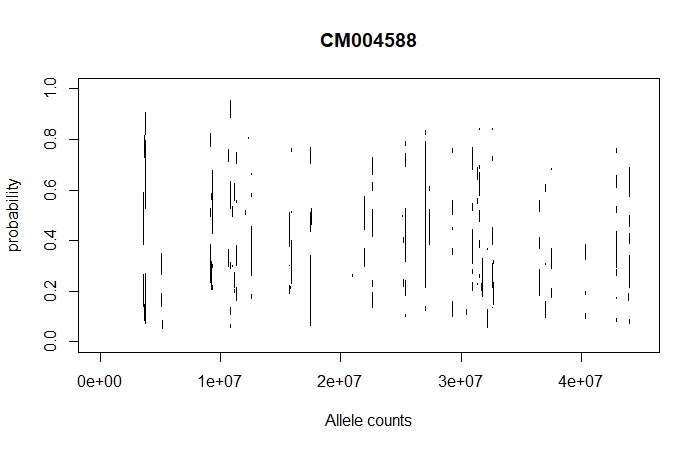

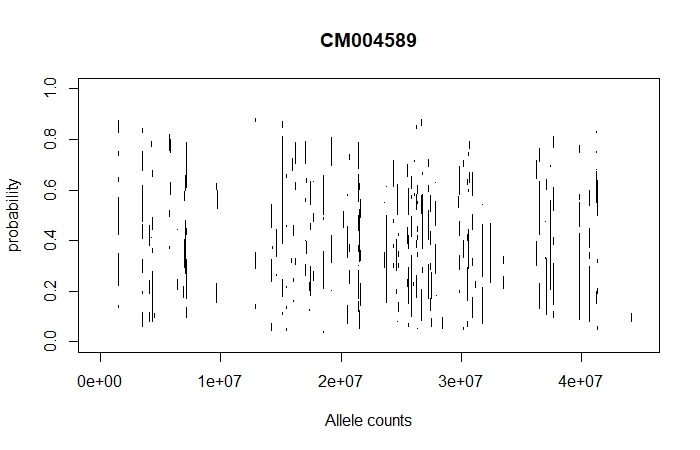

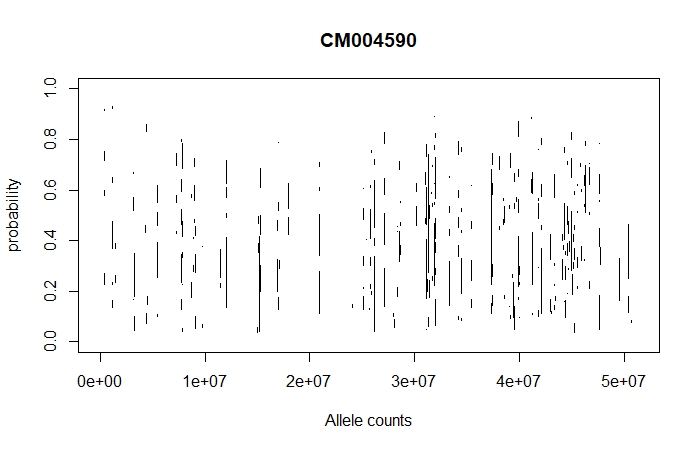

**Fig.S2** Graphical representation of allele counts and its probability across all autosomes of DDP goat.
